# RNAcomp2D: a web tool to execute multiple methods and visually compare predictions of RNA secondary structure

**DOI:** 10.1101/2025.10.08.681167

**Authors:** Rosario Vitale, Diego H. Milone, Georgina Stegmayer

**Affiliations:** Research Institute for Signals Systems and Computational Intelligence sinc(i) FICH-UNL CONICET Ciudad Universitaria UNL 3000 Santa Fe Argentina

## Abstract

Ribonucleic acids (RNAs) are involved in many important biological processes. In particular, non-coding RNAs are crucial regulators of cellular processes, playing a significant role in gene expression. RNA secondary structure is key to infer their specific function and for understanding how they interact with other molecules. Many computational models have been developed in the last decade to predict the secondary structure, achieving increasingly higher success rates. However, each new method has its own input-output interface, programming language, computational requirements and, sometimes, a dedicated server to run the model or just a source code in a repository. Thus, nowadays it is very hard to obtain predictions from multiple methods and compare them at once. A unified interface is urgently needed, which allows accessing several methods at the same time, visualizing and comparing predictions among them, and also with a reference structure when available. We introduce here RNAcomp2D, a web-based tool that allows users to enter an RNA sequence, or select one from RNAcentral, and obtain predictions of RNA secondary structures using several state-of-the-art methods. Both classical thermodynamic methods and the latest deep learning models are packaged in containers and accessible in an unified website. All the predictions, and the reference structure if available, are shown at the same time in a single graphical interface. Moreover, as new models continue to be developed, this tool is designed to be scalable, allowing the addition of more prediction methods in the future. Web server is available at https://sinc.unl.edu.ar/web-demo/rnacomp2d/. Data and source code are available at https://github.com/sinc-lab/RNAcomp2D

## 1. Introduction

For a long time, RNA was thought to act just as a messenger of information between DNA and proteins, playing a secondary role in the expression of information inherited from proteins. However, the recent ability to sequence genomes and transcriptomes has revealed that most of the human genome is transcribed, in fact almost 98% comprises non-coding RNA regions (1; 2). These regions, which are not used in protein synthesis, act on their own in diverse ways; thus knowing their secondary structure is the key for understanding their functions (3). The RNA molecule is an ordered sequence composed of four nucleotides or bases: adenine (A), cytosine (C), guanine (G), and uracil (U). The pairing of these four bases within an RNA molecule gives rise to its secondary structure (4; 5). These base pairs often result in a folded structure, where several pairs are stacked and unpaired bases form a loop. This secondary structure is usually represented in dot-bracket notation, with matching parentheses for paired bases and dots for unpaired bases. The most commonly used graphical representations of the folded structure are a 2D planar plot and a circular plot, which uses the clockwise order of the RNA sequence and arcs to indicate base pairs.

Many methods have been developed for the computational prediction of RNA secondary structure. The first proposals, dating back 25 years (6), were based on dynamic programming and thermodynamic calculations (7; 8), identifying a minimum free energy structure based on the principle that RNA molecules exist in energetically stable states (9). Until the emergence of machine learning-based methods in the field, about 15 years ago, and especially the deep learning (DL) ones in the last 5 years, the maximum possible accuracy to be achieved in computational predictions had remained almost unchanged (10). Most widely used classical methods include RNAFold (11), RNAstructure (12), LinearFold (13), LinearPartition (14), CONTRAfold (15), and IPKnot (16). DL models emerged as an alternative to classical approaches, making weaker assumptions about the thermodynamics driving RNA folding. There are many methods based on DL that appeared in the last 6 years (17; 18; 19), from the first proposals like SPOT-RNA (20; 21), to the most competitive up to date: UFold (22), REDFold (23), RNAformer (24), and sincFold (25). However, results from DL models evaluated on random partitions should be interpreted with caution, because performance may be overstated due to overfitting when training and testing data share sequences of the same RNA family (26).

Each DL model has its own architectural design, model input–output interface, computational requirements, training data and optimization algorithms used to adjust their parameters. Sometimes there is a dedicated server to run the model, but many times there is just a repository with instructions on how to implement it. To the best of our knowledge, there is no tool that allows running several methods concurrently, together with an integrated visual comparison of their computational predictions, from both classical and DL based methods, on a single interface. In spite of the existence of some secondary structure visualization libraries, those do not allow running different types of methods nor comparing many predictions at the same time on a single interface. For example, ViennaRNA (11) can compare only two secondary structures and RNAStructViz (27) can plot up to three structures simultaneously. Another example is RnaBench (**?**), which is a library specifically designed for the development of DL algorithms, providing datasets, evaluation metrics and a visualization module. It is a benchmark tool that helps mitigate the challenges of data generation, model evaluation, comparison and visualization of results. However, RnaBench is intended for benchmarking average predictors over entire datasets, and it was not conceived for web access, requiring local installation and not providing direct visual comparison of predictions, as RNAcomp2D offers. A very recent contribution is ShapeRNA (28), which despite running up to seven prediction methods, the predictions cannot be executed simultaneously requiring users to run each method individually. Furthermore, structure comparison remains a manual process as each prediction must be copied and pasted into a separate visualization module.

In this work we introduce RNAcomp2D, a web-based tool that allows users to enter an RNA sequence, or select one from the millions of sequences available at RNAcentral (29), and obtain its corresponding secondary structure using several existing state-of-the-art methods, both classical and DL based. All the predictions, and the reference structure if available, are shown in the same graphical interface and can be downloaded as well in several formats. RNAcomp2D as a web application does not require installation, it can be freely accessed with any web browser, and it allows a detailed visual comparison of the predicted secondary structure for one sequence at a time, with many prediction methods (up to eleven in the current version). The similarities and differences among all the predictions are indicated with different colors and intensity grades in the circular plot. The main aim of this tool is visualization, for helping both biologists and computational scientists developing new methods, to graphically compare the predictions of existing methods and the reference structure.

## 2. The RNAcomp2D web tool

The web interface allows the user to enter a single RNA sequence to be folded: i) manually, by copy and paste of a sequence in the text box; ii) by uploading a FASTA file; or ii) by retrieving the sequence from RNAcentral^1^. After entering the RNA sequence, the user can select one or more prediction methods to be compared. Then, the request is submitted to the web server. The resulting secondary structures predicted are then generated and displayed as a graphical planar plot and a circular plot. In the case of retrieving a sequence from RNAcentral where the corresponding reference structure is available, or if the user uploads a FASTA that includes a reference structure, it will be shown first and remain pinned in the results page, while the predicted structures will be shown at the right in order to be compared to the reference. The *F*_1_ score, calculated between the reference structure and each prediction method output, is also provided. These steps are explained in more detail in the following subsections.

### 2.1. Sequence input and RNAcentral search

There are several options to input a sequence into the RNAcomp2D visual interface. In the main text box (Figure 1, left panel), the RNA sequence can be entered either by typing nucleotides (or copy-paste them), or by clicking on one of the examples provided. Below this box, there is a button to upload a file in FASTA format. The file must contain the sequence, and may also include a reference structure in dot-bracket notation. To indicate the reference structure, the identifier must contain the word “Reference”. Additionally, in the same file, predictions from other methods can be included as well to be compared graphically.

**Figure 1:**
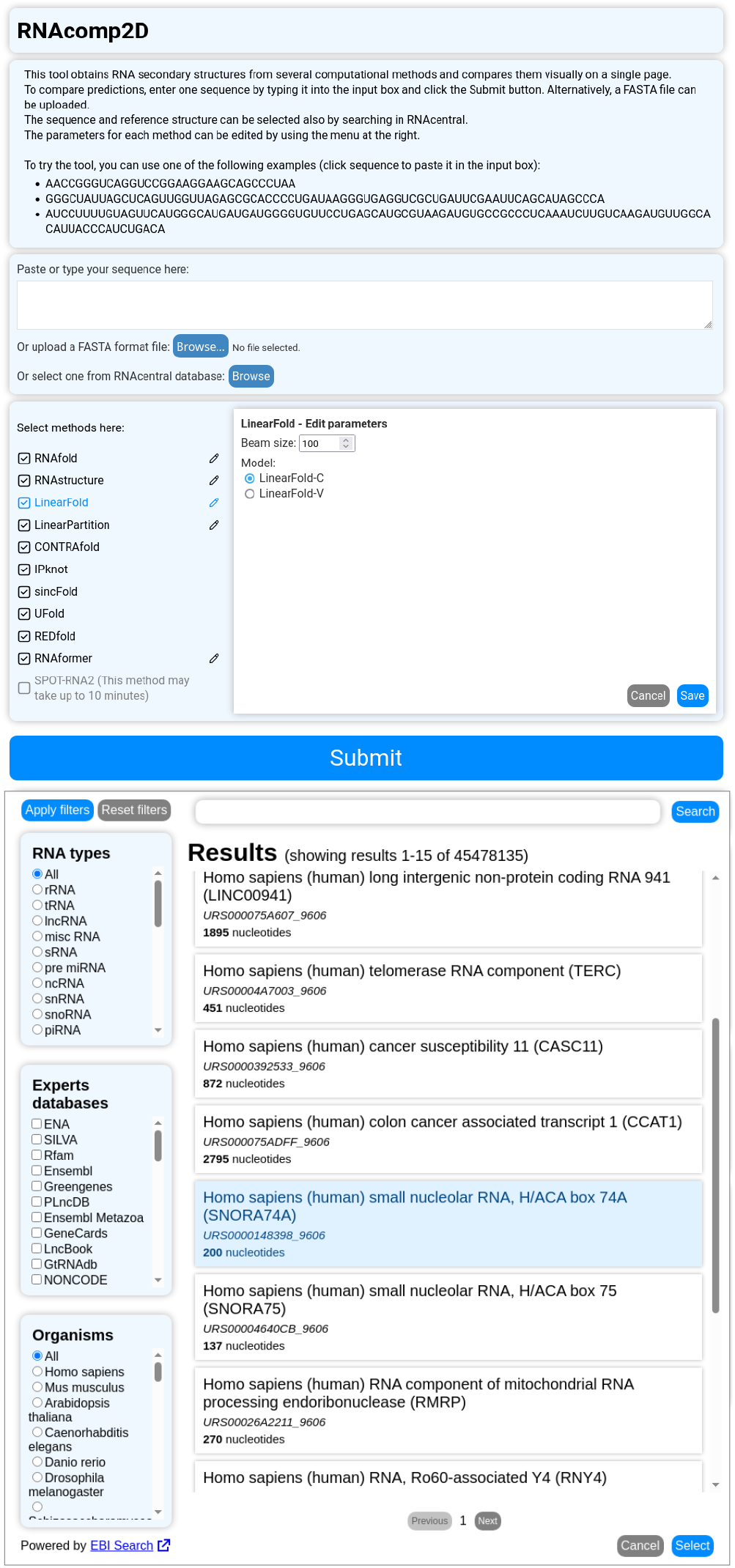
Main window of RNAcomp2D that allows entering the RNA sequence to be folded (left panel). Window to retrieve a RNA sequence from RNAcentral (right panel).

There is also a Browse button to search a sequence via RNAcentral (29). It is a public database hosted by the EBI Institute (30) that offers access to a complete and up-to-date set of more than 30 million RNA sequences. It is updated twice a year, each time resulting in a new version that incorporates and revises the available sequences (31). PDB sequences are also included in RNACentral, together with the reference RNA secondary structure, when available. Retrieving any RNA sequence from RNAcentral and then comparing several prediction methods for this sequence is a very important feature of RNAcomp2D. At the dialog box for RNAcentral (Figure 1, right panel) users can enter a search term, e.g. *hsa-miR-1-5p*, and can also refine the results by using several available filters at the left. The filters allow searching by RNA types, expert databases and by organisms. Once the desired sequence is found, users can select it and close the dialog. If a reference structure is available for the selected RNA entry in RNAcentral, it will be retrieved and used for visual comparison with the computational predicted structures and for the calculation of the *F*_1_ score.

In the case of methods that require parameters, the values provided by original authors are used by default, which however can be changed if necessary in the main interface of RNAcomp2D. For RNAfold, temperature, salt concentration and the option to produce structures without lonely pairs can be set. For RNAstructure, temperature and maximum loop size can be changed. For LinearFold, the beam size and the option to use the machine learned scoring model (LinearFold-C) can be chosen. In LinearPartition, the last option for machine learning derived parameters can be selected as well. For RNAfomer, the cycle parameter (number of model iterations) can be edited, and the available models can be selected (biophysical, bpRNA, intra family finetuned or inter family finetuned). There is a practical limitation for users of some DL methods regarding the maximum sequence length allowed for predictions. For sincFold and REDfold it is 512 nt, while for UFold it is 720 nt.

### 2.2. Selection of RNA secondary structure prediction methods to compare

To compare predictions of several methods among them, below the sequence entry section there is the method selection section, which consists of a list of available methods showing which ones will be executed once the request is sent to the server. By default, all methods are selected. To the right of methods, an edit icon allows configuring parameters (see example in Figure 1, left panel). In some cases, these parameters can significantly alter the results obtained, such as the temperature in the case of thermodynamic methods.

The main selection criteria for including prediction methods in RNAcomp2D is to provide access to the most highly cited classical methods and most competitive DL based methods, according to recent literature reviews (**? ? ? ? ?** 26; 19). An additional criteria for methods inclusion is their availability as installable packages or as free source code available in public repositories. Therefore, the classical methods RNAfold 2.7.0, RNAstructure 6.5, LinearFold 1.0.1, LinearPartition 1.0, CONTRAfold 2.02 and IPknot 2.3.1 are included, together with the DL based methods SPOT-RNA2, UFold 1.3, REDFold 1.14, RNAformer 0.0.1 and sincFold 0.16.2. For each prediction method, we followed the instructions available in their repositories. The thermodynamic methods were compiled from the source codes, while the DL methods were installed and the corresponding pre-trained models were downloaded from repositories. Each method runs with its own thread at the web server. Docker containers (32) were built allowing better handling of computational requirements at the server side and future extensions to other prediction methods.

### 2.3. Visual comparison of predictions and performance analysis

Once the RNAcomp2D server receives the request, all the selected prediction methods are executed and the corresponding plots are generated for each predicted secondary structure. Prediction results are displayed on screen as they become available. For longer sequences, some methods may require several minutes to finish. Figure 2.A, shows a screenshot of the RNAcomp2D results page. The top bar allows the user to change the size of the displayed images, select the type of output representation and download all method predictions. The Download button opens a dialog to select which results to download (figures in PDF or predictions in FASTA). At the bottom of the page, the identifier of the RNA (in this example, the tRNA URS000007A90A_9606 from RNAcentral) and its corresponding sequence can be seen.

**Figure 2:**
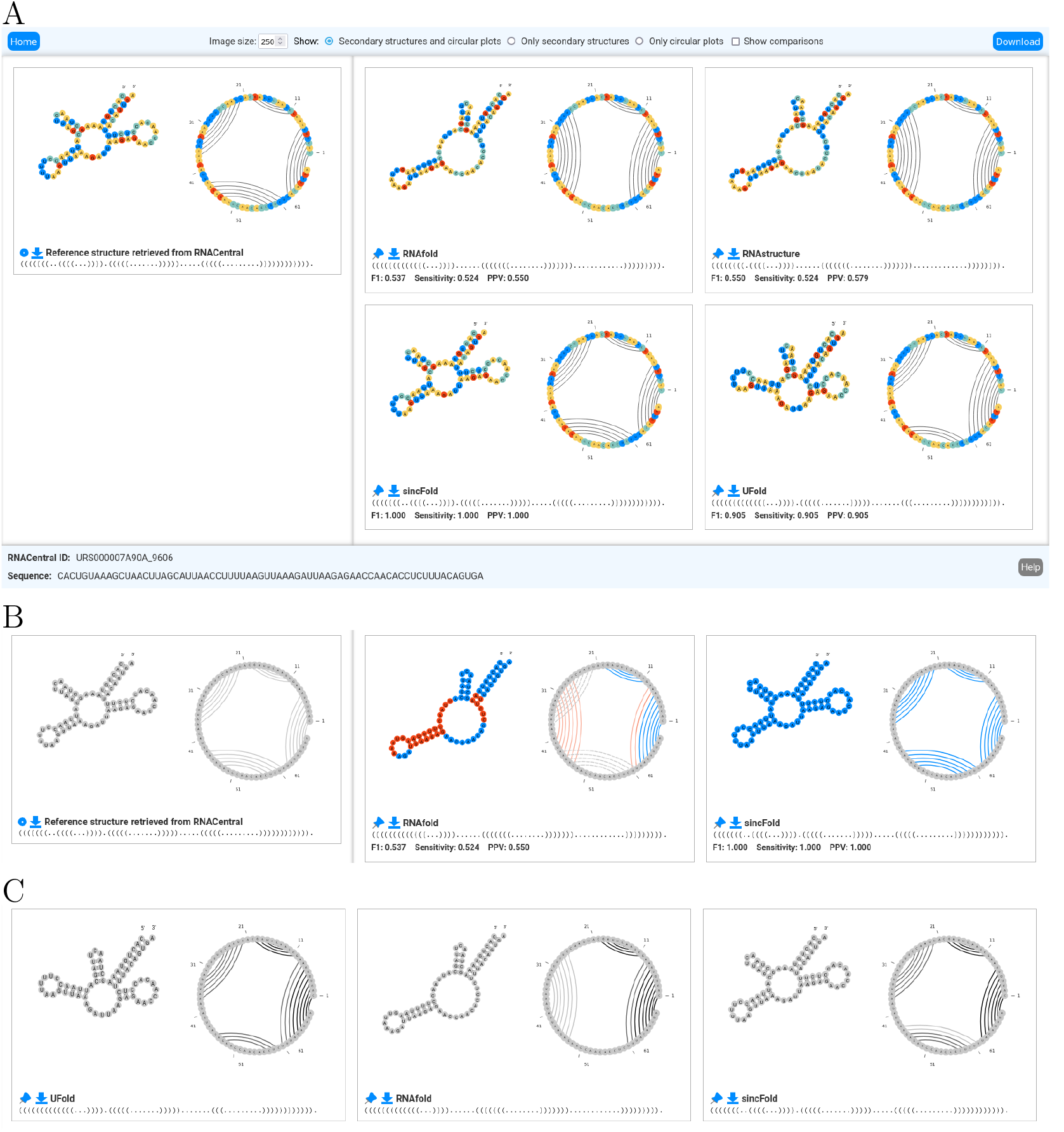
RNAcomp2D results page for the visual comparison of computational methods for RNA secondary structure prediction. A) Screenshot of the default page of results, with a reference structure pinned at the left. B) Output screen for the “Show comparisons” option when a reference structure is available: base pairs and connections predicted correctly, incorrectly or not predicted are shown with different colors. C) Output screen for the “Show comparisons” option when there is no reference structure: gray scale indicates the amount of predictions shared among methods.

The prediction results are displayed in the form of a grid of images, one for each prediction method. Each cell in the grid indicates the name of the method and the prediction is shown according to the output graphical representations selected: plane 2D structure^2^ and circular plot. Each base type is indicated with a different color, and it should be mentioned that the color palette has been selected to be suitable to colorblind users. Users can download the individual prediction of each method using the download button on the left of the method name. Clicking this button opens a dialog where the user can choose to download the dot-bracket prediction in FASTA format, or its graphical representation in PDF, SVG, PNG, or JPG format. If a method is pinned, its cell is fixed on the top-left of the page. If a reference structure exists (in RNAcentral or it is provided by the user) it is always pinned by default on the top-left of the web page.

When there is a reference secondary structure, the sensitivity, positive predictive value (PPV), and *F*_1_ score are calculated for each prediction, providing metrics that quantify the similarity between the predicted base pairs and those in the reference structure. The base pairs that are both in the prediction and in the reference structure are true positives (TP). The pairs that are only in the predicted structure are false positives (FP). A pair in the reference structure that is not predicted is a false negative (FN). Then, the *F*_1_ score is computed as 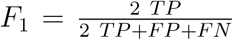. The PPV measures the precision of predicted base pairs calculated as the fraction of predicted pairs that exist in the reference secondary structure, that is 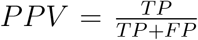. Finally, sensitivity or recall is calculated as 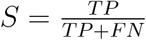.

The results interface also provides a “show comparisons” option, allowing the comparison of all the predicted RNA structures among them, indicating the shared connections. When there is a reference structure, the base pairs that were predicted correctly, incorrectly or not predicted by the methods are shown with different colors in both the secondary structure representation and the circular plots (blue for TP and red for FP, see Figure 2.B). Additionally, the color intensity in the circular plot is proportional to the number of methods that share a predicted connection. When there is no reference structure, a gray scale is used to indicate the amount of predictions shared (Figure 2.C). This visualization allows easily see, at a glimpse, which base pairs and connections are more trusted by the majority of methods.

Regarding runtime of methods, for a sequence of 100 nt, the approximate execution time for each classical method is between 3-4 s, while for DL models it is between 12-18 s. Only for SPOT-RNA2 the execution times are significantly longer, around 1000 s on average, mainly due to the computational overhead of BLAST preprocessing. A detailed runtime analysis of the prediction methods included, across a range of sequence lengths from 33 to 2,000 nt, is depicted in Supplementary Figure S1. The execution time increases almost linearly with sequence length in the case of DL methods. With the exception of SPOT-RNA2, DL methods share a lower bound of approximately 10 s in prediction times, suggesting that model instantiation and parameter loading dominates runtime for short to medium sequences. SPOT-RNA2 consistently presents large prediction times because it has an additional fixed bottleneck arising from a remote BLAST call prior to folding. In contrast, classical methods exhibit a wider range of scaling behaviors, from near-linear growth (e.g., LinearFold) to the cubic time complexity theoretically expected for these methods, and even higher-order polynomial growth (e.g. IPKnot). The use of memory by each method is largely variable (Supplementary Table S1), with thermodynamic approaches requiring approximately 250 MB, while DL methods demand considerably more, ranging from 860 MB (REDfold) to 6.80 GB (RNAformer). The web server has an Intel Xeon Gold processor 5520+, 64 GB of RAM and a HD of 3.84 TB. The virtual machine of RNAcomp2D has been assigned 7 cores, 10 GB of RAM and 50 GB of HD.

A key advantage of RNAcomp2D is its asynchronous architecture: each method runs in a dedicated execution thread at the server. Therefore, each prediction is processed in parallel and is shown at the output interface as soon as it arrives at the client side. Regarding queues and concurrency behavior, for each job submission the selected prediction methods are launched in independent execution threads, each invoking a separate prediction process that allows them to run in parallel. The server assigns a unique session identifier to each request and creates a dedicated working directory where all intermediate and output files are stored, preventing conflicts among concurrent executions. RNAcomp2D does not implement an explicit job queue or enforce application-level limits on the number of simultaneous submissions or running methods. Instead, concurrent execution is managed by the operating system scheduler, which allocates the available CPU and memory resources among active processes. The timeout of failure policy we have implemented allows each session to remain active for up to 1 hour, after which the resources are released. During our evaluations, all tested prediction methods completed within this time limit. If a prediction method terminates with an error, its execution thread is stopped and the corresponding error message is recorded and displayed on the results page, allowing users to identify failed methods while preserving the results of successfully completed ones. No failures due to resource exhaustion were observed during testing. Server resource utilization is continuously monitored, and automated alerts are generated when sustained CPU or memory usage exceeds predefined thresholds, to facilitate timely administrative intervention.

## 3. Conclusions

In this work we presented RNAcomp2D, an integrated web tool for the computation and visual comparison of secondary structures of RNA predictions. It integrates eleven state-of-the-art methods through an unified web interface: six based on thermodynamic models, and five on the most recent deep learning approaches. While the rapid evolution of prediction algorithms often renders tools obsolete, RNAcomp2D specifically mitigates this limitation by packaging methods or groups of similar methods in Docker containers for easy setup and extension. Predictions from all methods are graphically presented on a single screen, allowing a direct analysis and visual comparison. The user can upload an RNA sequence or select one RNA from the millions available at RNAcentral, with several filters to facilitate the search. All prediction methods are executed concurrently in parallel threads, and results are rendered progressively as they arrive from the server side, substantially reducing response times. When the reference secondary structure is available, either retrieved from RNAcentral or provided by the user, it is shown pinned and all the computational secondary structure predictions are compared visually, including also *F*_1_ score, precision and sensitivity. Furthermore, even without a reference structure, the interface provides a visual comparison of all the structures predicted indicating the shared connections with different color intensities proportional to the number of methods that agree on each pairing. In summary, RNAcomp2D provides a scalable framework for prediction and comparative visualization of RNA secondary structures, designed to be scalable and adaptable to new methods emerging in the field.

## 4. Acknowledgments

Authors thank Agustin Baricalla and Prof. Uciel Chorosteki for their useful revisions and comments to improve this tool. Authors also thank Prof. Matias Gerard for providing the initial implementation of the circular plot.

## 6. Author contributions statement

G.S. and D.H.M.: Conceptualization, Supervision, Formal analysis and Funding acquisition. R.V.: Data curation and Software development. D.H.M.: Visualization. G.S.: Writing – original draft. All authors: Writing – review & editing.

## Footnotes

1 https://www.rnacentral.org/

2 Generated with the draw_rna library: https://github.com/eternagame/draw_rna

