## Supplementary material for "RNAcomp2D: a web tool to execute multiple methods and visually compare predictions of RNA secondary structure"

Research Institute for Signals, Systems and Computational Intelligence, sinc(i), FICH-UNL, CONICET, CCT-Santa Fe, Ruta Nacional N° 168 km 0, Paraje el Pozo, Santa Fe, Argentina.

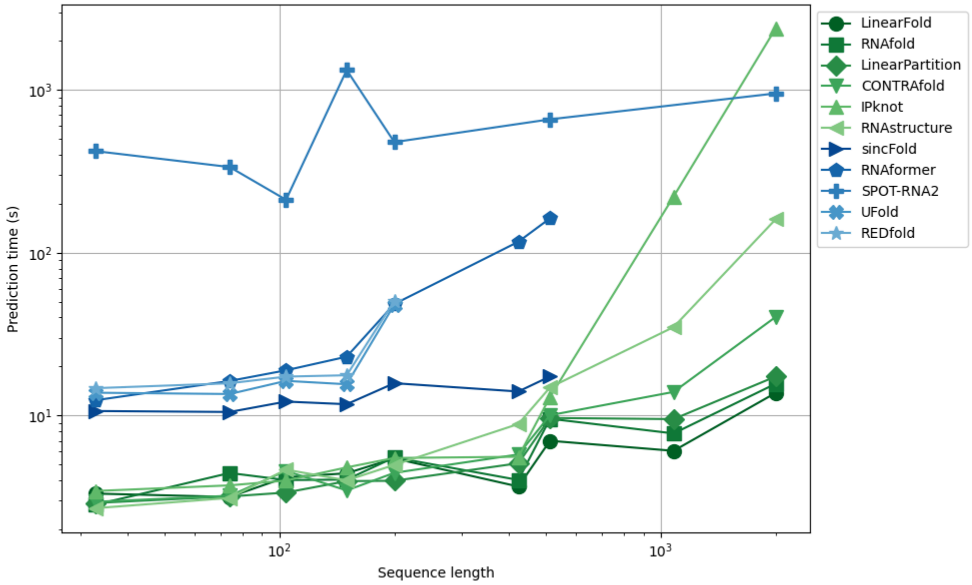


**Figure S1**. Prediction runtimes for methods included in RNAcomp2D: sequence length (x-axis) versus prediction time in seconds (y-axis). UFold, sincFold, and REDfold have been evaluated up to a specific sequence length because of model restrictions.

**Table S1**. Detailed memory use of prediction methods for a RNA sequence of length 512 nt.

| Prediction method | Memory use |
|---|---:|
| RNAfold | 258.52 MB |
| RNAstructure | 225.03 MB |
| LinearFold | 258.89 MB |
| LinearPartition | 250.94 MB |
| ContraFold | 264.20 MB |
| IPknot | 269.20 MB |
| REDfold | 860.49 MB |
| UFold | 984.15 MB |
| SPOT-RNA2 | 1.28 GB |
| sincFold | 1.45 GB |
| RNAformer | 6.80 GB |
| All methods | 7.61 GB |
